# Targeting the Tumor-Stroma Crosstalk: An AI-Based Virtual Screening Strategy for Dual MET/SMO Inhibitors in Pancreatic Cancer

**DOI:** 10.64898/2026.07.03.736313

**Authors:** Michele Roggia, Ugo Chianese, Giorgio Amendola, Valentina Albanese, Cinzia Vetrei, Caterina Ieranò, Crescenzo D’Alterio, Salvatore Di Maro, Fortunato Ciardiello, Floriana Morgillo, Stefania Scala, Lucia Altucci, Delia Preti, Gunnar Schulte, Rosaria Benedetti, Pawel Kozielewicz, Sandro Cosconati

## Abstract

Pancreatic ductal adenocarcinoma (PDAC) is an aggressive malignancy characterized by a dense desmoplastic tumor microenvironment (TME) that limits drug penetration and promotes immune evasion. Effective treatment, therefore, requires simultaneous modulation of multiple signaling pathways. Here, we describe a directed polypharmacological strategy to identify dual modulators of c-MET and Smoothened (SMO), aiming to disrupt the protective stroma through SMO inhibition while directly suppressing tumor cell survival via c-MET targeting.

An AI-guided virtual screening workflow combining the machine-learning platform PyRMD, trained on known c-MET and SMO ligands, with structure-based molecular docking was applied to a library of over 9 million compounds. This approach led to the identification of compound **21**, an aminopyrimidine-benzamide-phenoxyquinoline derivative, as a dual c-MET/SMO inhibitor.

Biochemical and cellular studies demonstrated that compound **21** selectively binds the SMO orthosteric site (pKi = 5.60), inhibits agonist-induced GLI (Glioma-associated oncogene) signaling (pIC_50_ = 5.50), and potently suppresses c-MET kinase activity (pIC_50_ = 6.94). Western blot analyses further revealed that compound **21** promotes ubiquitin–proteasome-mediated degradation of c-MET, eliminating receptor availability and limiting compensatory resistance signaling. In 3D heterotypic models comprising MIAPaCa2 pancreatic cancer cells and CAF154-hTERT fibroblasts, dual inhibition of SMO-mediated stromal support and c-MET-driven tumor progression resulted in greater cytotoxicity than the combination of the selective inhibitors Sonidegib and PHA-665752. Overall, compound **21** overcomes stromal-mediated resistance, enhances tumor cell death, and validates dual SMO/c-MET targeting as a promising single-agent therapeutic strategy for PDAC.

**One Sentence Summary:** An AI-identified dual SMO/c-MET inhibitor overcomes stromal resistance and degrades c-MET to suppress pancreatic cancer.

## INTRODUCTION

Pancreatic ductal adenocarcinoma (PDAC) is an aggressive malignancy characterized by dismal prognosis and limited treatment options (*1*). It remains one of the most lethal cancers, with a 5-year survival rate of approximately 10%, a scenario that has seen only modest improvement despite advances in oncology(*2*). The prognosis remains severe even after complete surgical resection due to high recurrence rates (*3*). Current therapies have a minimal impact on the disease, with the long-term survival rate barely exceeding 20%(*4, 5*).

The current standard regimens include FOLFIRINOX (5-fluorouracil, leucovorin, irinotecan, and oxaliplatin) and the combination of gemcitabine with nab-paclitaxel, though both offer modest survival benefits(*6*). Treatment efficacy is frequently hindered by the desmoplastic and immunosuppressive tumor microenvironment (TME), which also renders PDAC largely unresponsive to immune checkpoint inhibitors (ICIs)(*7*). Resistance to conventional and immunotherapeutic strategies underscores the urgent need for innovative approaches targeting the molecular drivers of PDAC progression. Immune resistance is primarily ascribed to the deeply immunosuppressive tumor microenvironment, characterized by dense stroma, low tumor mutational burden, and limited effector T-cell infiltration (*8*). Consequently, immunotherapy has failed to produce practice-changing results in PDAC, with responses largely restricted to the rare subset of patients harboring mismatch repair deficiency or microsatellite instability-high tumors (*9*).

Multifactorial diseases like cancer may benefit from polypharmacology, the design of drugs capable of targeting multiple oncogenic pathways simultaneously. Unlike single-target therapies, polypharmacological agents can disrupt multiple signalling networks, enhancing efficacy and overcoming resistance mechanisms (*10*). The clinical rationale relies on data showing that combination therapies targeting distinct molecular pathways often yield synergistic effects, improving patient outcomes beyond the sum of individual drug benefits (*11*). Nonetheless, compared to combination therapy, polypharmacological agents offer several advantages, including simplified pharmacokinetics, the absence of drug-drug interactions, and improved patient compliance (*12*). Most importantly, they can simultaneously disrupt compensatory signalling pathways, thereby more effectively preventing the emergence of drug resistance (*10*). Our previous research identified dual inhibitors of the receptor tyrosine kinase c-MET (Hepatocyte growth factor receptor) and the Smoothened (SMO) Hedgehog (HH) pathway, which are implicated in resistance to EGFR inhibitors in lung cancers (NSCLC)(*13*). The rationale for targeting these targets in NSCLC stemmed from their role in promoting epithelial-to-mesenchymal transition (EMT), a key process in tumor progression and metastasis(*14*).

Given that NSCLC and PDAC share aggressive features(*15*), we hypothesized that simultaneous targeting of c-MET and SMO may also be beneficial in PDAC patients. Recently, PD-1 cancer signalling was described in PDAC (*16*). While classically recognized as an immune checkpoint receptor that induces T-cell exhaustion, PD-1 is also endogenously expressed on pancreatic ductal adenocarcinoma (PDAC) cells, where it assumes a distinct oncogenic role. Contrary to its established function in immune regulation, tumor-endogenous PD-1 activates the proto-oncogene c-MET. Mechanistically, stimulation of tumor-intrinsic PD-1 does not involve direct protein-protein interaction with c-MET but rather triggers the transcriptional upregulation of the c-MET ligand, the Hepatocyte Growth Factor (HGF). HGF subsequently activates c-MET via autocrine signalling, driving the downstream epithelial-to-mesenchymal transition (EMT)(*17*). This axis, operating independently of immune cell interactions, enhances cancer cell growth, migration, and invasion, rendering PDAC cells more aggressive through epithelial-to-mesenchymal (EMT), a well-established early oncogenic process in PDAC (*18*). Thus, the inhibition of c-MET can reverse the EMT phenotype in PDAC.

Hedgehog (HH) pathway is a crucial driver of the dense desmoplastic stroma characteristic of PDAC. In this paracrine signalling loop, neoplastic epithelial cells secrete Sonic Hedgehog (SHH), which binds to the receptor Patched-1 (PTCH1) on adjacent cancer-associated fibroblasts (CAFs). HH binding to PATCH1 relieves the repression of the transmembrane protein Smoothened (SMO), triggering a cholesterol-driven cascade that activates GLI transcription factors and drives excessive extracellular matrix deposition. The resulting dense stroma acts as a formidable physical barrier, creating a hypoxic environment that severely restricts chemotherapeutic delivery. Consequently, inhibiting the HH pathway via SMO antagonists or inverse agonists has shown promise in ‘priming’ PDAC tumors by reducing stromal density and improving drug penetration (*19*)(*20*). Surprisingly, accumulating evidence also suggests that disrupting HH signalling, particularly through prolonged SMO inhibition, can paradoxically promote EMT in PDAC cells. This occurs as HH pathway blockade releases stromal constraints on tumor cells, fostering de-differentiation and a more invasive, mesenchymal phenotype (*21–23*). Furthermore, recent studies have illuminated the intricate role of the inflammatory TME in modulating HH pathway activity in PDAC. Pro-inflammatory cytokines, such as TNF-α and IL-1β, abundantly present in the PDAC stroma, activate the HH pathway in PDAC cells, promoting EMT and other malignant behaviors(*24*). Thus, mounting evidence suggests that, similar to observations in NSCLC, SMO and c-MET represent highly relevant targets for polypharmacological intervention in PDAC. While SMO inhibition provides a therapeutic ‘priming’ effect by reducing stromal density and improving tumor access, it also carries the inherent risk of triggering a compensatory EMT, which can lead to increased invasiveness. By concurrently inhibiting c-MET, this pro-metastatic shift is effectively neutralized. Therefore, a dual-inhibitor approach, targeting SMO for its priming capabilities alongside c-MET to counteract EMT, could be particularly valuable in optimizing therapeutic outcomes for PDAC. Designing a single agent capable of binding and modulating multiple pharmacological targets presents a significant challenge. This is due to the complex interplay of factors arising from the distinct structural demands of each target, including variations in binding site architecture and molecular recognition elements (*25*). Accommodating the diverse shapes, sizes, and chemical properties of multiple binding pockets within a single compound is a major hurdle, especially considering that, in our case, c-MET and SMO are phylogenetically distinct biological targets. Furthermore, achieving the necessary specificity and selectivity to ensure binding to the desired targets while avoiding off-target interactions is critical. In this scenario, the application of a computational design strategy is desired. If this strategy takes into consideration the structural demands that control how a drug-like molecule and its target recognize one another, the strategy could allow the detection of shared pharmacophoric features between the targets of interest (*10*).

Herein, we implemented a customized virtual screening (VS) protocol using PyRMD(*26*), an AI-powered ligand-based VS tool designed to facilitate the rapid and accurate identification of novel biologically active compounds. To address the structural diversity of the kinase target, we tailored the protocol by generating distinct prediction models for Type I and Type II c-MET inhibitors, effectively stratifying the screening of the Mcule database (∼9 million compounds) based on specific binding modes. The resulting hits were subsequently cross-referenced with a SMO-specific model and subjected to rigorous structure-based filtering, which prioritized candidates capable of interacting with critical residues in both the c-MET hinge region – the ATP binding site - and the SMO transmembrane domain. This approach successfully identified compound **21**, a unique single-agent polypharmacological modulator. Our results demonstrate that compound **21** blocks SMO signalling, selectively inhibits c-MET activity, and inhibits cell proliferation of pancreatic cells while sparing non-cancerous pancreas cells. Furthermore, in complex 3D heterotypic spheroids, this dual-targeting strategy exhibited superior cytotoxicity compared to the combination of selective single-target inhibitors, successfully overcoming the structural resilience of the tumor stroma. This polypharmacological approach represents a step toward more effective treatment strategies by targeting key oncogenic pathways involved in PDAC progression and resistance mechanisms.

## RESULTS

### AI-enforced Virtual Screening

To search for dual c-MET/SMO binders, a hybrid approach was adopted (Fig. 1). Firstly, a ligand-based screening strategy for the c-MET target, powered by the AI-based PyRMD software, was employed. PyRMD’s machine learning algorithm, trained with ChEMBL data, an open-access, manually curated database of bioactive molecules with drug-like properties, allows for the prediction of potential binders against a specified target. The software’s data processing module automates the separation of the ChEMBL database into active and inactive compounds. These sets were then used to develop a machine learning model capable of differentiating structural features between biologically active and inactive compounds.

**Fig. 1.**
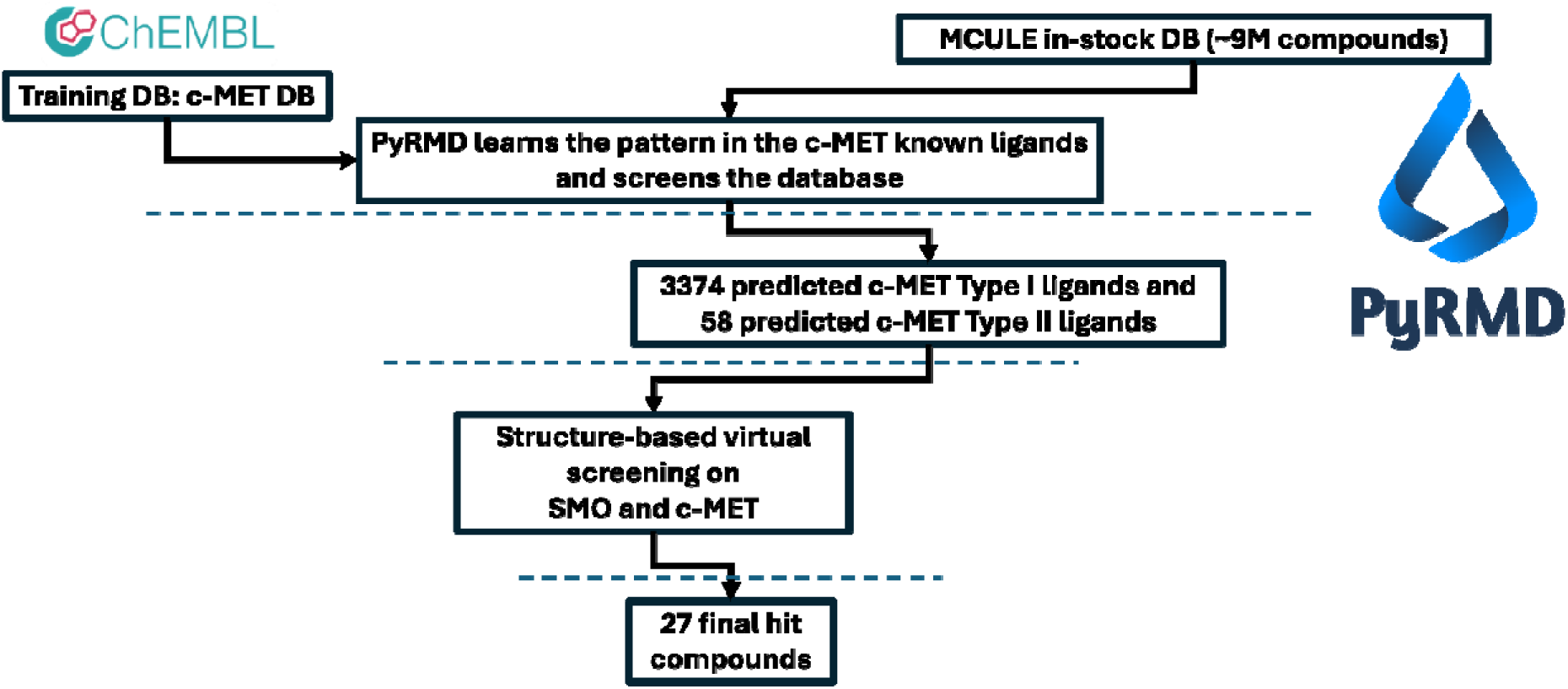
AI-driven hybrid virtual screening workflow for the discovery of dual c-MET/SMO modulators. The computational pipeline integrates ligand-based machine learning with cross-target molecular docking to isolate dual-target hits from over 9 million in-stock compounds. First, PyRMD machine learning models were trained on ChEMBL data to specifically recognize the geometric features of Type I and Type II c-MET inhibitors, rapidly narrowing the Mcule database to a focused subset of candidate hits. To establish dual-target affinity, these candidates were then advanced to structure-based molecular docking against an ensemble of c-MET and SMO crystal structures. Final selection prioritized compounds capable of simultaneously engaging critical residues in both the c-MET ATP-binding hinge region and the SMO orthosteric pocket, culminating in a final set of testable hits for biological validation.

Therefore, the database of the reported bioactivities associated with the human c-MET (9928 entries) was downloaded from ChEMBL. Here, different types of c-MET inhibitors have been reported with distinct chemical features that interact with different portions of the ATP-binding pocket of c-MET. Therefore, we decided to create two separate PyRMD models for the two main c-MET inhibitor categories (i.e., type I and type II). Type I inhibitors possess a U-shaped binding mode while type II inhibitors extend through the gatekeeper portion of the binding site, occupying the deep hydrophobic back pocket of the catalytic site(*27*). Thus, all compounds downloaded from ChEMBL were classified into two sets, or discarded if a clear decision could not be made, by evaluating two distinct criteria: the shape overlap with two c-MET inhibitor that prototypically represent type I and type II classes (PDB: 4R1V(*28*) and PDB: 4MXC(*29*), respectively, selected for their high crystal resolution), and the presence of hydrogen bond acceptors and donors essential for interacting with the c-MET hinge region. To this aim, we employed the shape screen module available in Maestro within the Schrödinger suite(*30*). Next, the two sets were used as training data for the PyRMD benchmark module, where different settings (e.g., activity thresholds, ε cut-off values) were tested to identify the best-performing combination. The parameters that yielded the best results in terms of F-Score were selected for the type I model and the type II model. The two different PyRMD models were then used to screen the Mcule database of in-stock compounds (∼9 million compounds). The type I model returned 3374 possible inhibitors, while the type II model returned 58 compounds. The construction of a robust ML-based prediction model via PyRMD requires setting widely separated bioactivity thresholds to guarantee an accurate separation between active and inactive datasets, as implemented for c-MET. Unfortunately, applying such a stringent separation to the SMO dataset led to a significant depletion of available data points. This data scarcity severely impaired the model’s training process, precluding effective learning and ultimately yielding a high number of false positives. Thus, to address this challenge and ensure the successful identification of dual c-MET/SMO modulators, the compounds prioritized as actives from the c-MET screening were directly subjected to a classical structure-based virtual screening (SBVS) campaign. Specifically, molecular docking simulations against SMO were carried out using the methodology optimized and validated in our previous work employing six different SMO crystal structures (i.e., 4JKV, 4N4W, 4O9R, 4QIM, 5L7I, 5V57). In this context, we retained only compounds whose docking pose interacted with receptor residues critical for binding (N219, W281^2.58^, D384, Y394, R400^5.43^, F484^6.65^, and E518^7.38^)(*31*).

To further refine our results, we also performed molecular docking studies on the c-MET type I and type II crystal structures (PDB codes 4R1V, 4MXC, respectively). In this analysis, for both c-MET types, we selected only compounds predicted to interact with the hinge region. Compounds that met the criteria for both targets were clustered and, through final visual inspection, a set of 39 molecules was selected. Encouraged by this evaluation, we decided to purchase the 27 commercially available molecules as pure substances and test them. During testing, 4 of these 27 molecules were insoluble and therefore discarded.

### The newly identified candidates are SMO antagonists

The 23 compounds for which a dual SMO/c-MET binding behaviour was predicted were first screened for their ability to bind the SMO receptor with a NanoBRET-based ligand competition binding assay(*32*). In the first screening experiment, all the compounds were used at 10 μM and were tested for their ability to compete out the fluorescently tagged SMO antagonist cyclopamine (BODIPY-cyclopamine) used at 10 nM. As the compounds are predicted to target the orthosteric binding site of the receptor, N-terminally Nluc-tagged SMO lacking the CRD domain was used (ΔCRD Nluc-SMO), as this has been shown to increase the window of the binding assay. According to the results shown in Fig. 2A, compounds **2**, **6**, **21,** and **22** visibly decreased the BRET (approximately to the −3xSD, green dashed line), suggesting competition with the fluorescent probe (black dashed line indicates BRET in the absence of any competing ligand), indicating that the compounds bind to the BODIPY-cyclopamine binding site in the core of the receptor. Based on the initial results, these four compounds were used in the competition binding assays employing a full concentration range for each of them. Here, compound **21** showed the highest affinity (pKi=5.6 ± 0.4) while the other compounds bound to SMO with lower affinities (**2**: pKi=5.0 ± 0.4; **6**: pKi=3.5 ± 1.7; **22**: pKi=5.0 ± 0.1) (Fig 2B). Next, in order to assay the mode of action of compound **21**, we performed a GLI reporter assay as a functional signalling readout. To investigate the inhibition of SMO-mediated signalling by compound **21**, the Shh Light II cell line, stably expressing a GLI-responsive Luciferase reporter, which produces luminescence when the Shh pathway is activated, was used. Here, we assessed the ability of compound **21** to affect agonist (SAG21k)-induced, SMO-dependent GLI signalling. Indeed, compound **21** blocked SMO agonist-induced signalling in Shh-Light II stably overexpressing the GLI reporter with a pIC_50_=5.5 ± 0.0 (Fig 2C).

**Fig. 2.**
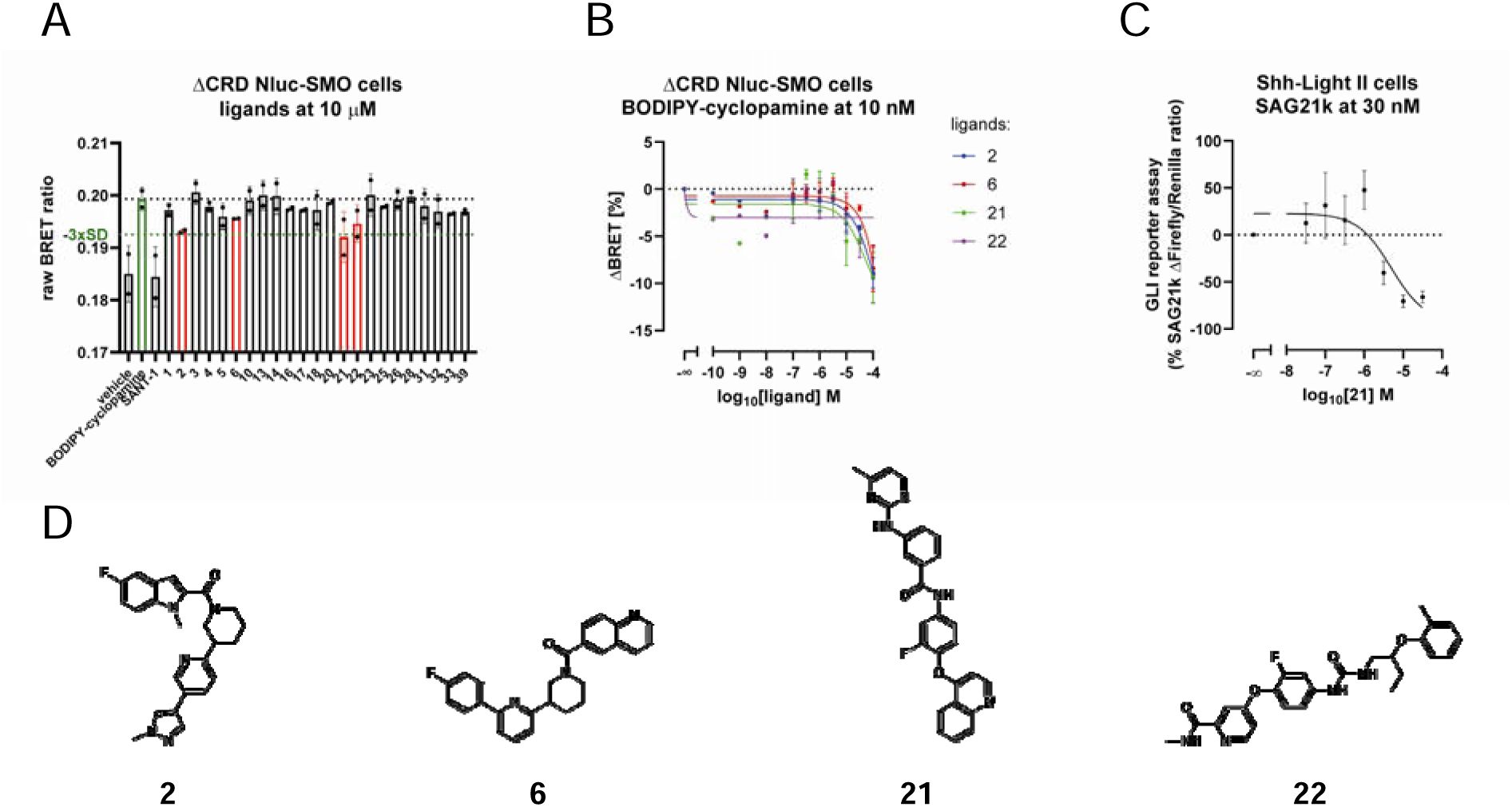
A) Compound screen using a direct binding assay in HEK293A cells stably expressing ΔCRD Nluc-SMO. Cells were preincubated with 10 μM of the compounds for 30 min, and subsequently 5 nM of BODIPY-cyclopamine were added and the cells incubated for an additional 90 min. The data are presented as mean ± SD. of two independent experiments, each performed in technical triplicate. B) Competition ligand binding assay in HEK293A cells stably expressing ΔCRD Nluc-SMO. Cells were preincubated with a full concentration range of **2**, **6**, **21**, **22** and subsequently 10 nM of BODIPY-cyclopamine were added, and the cells were incubated for additional 90 min. The data are presented as mean ± SEM of three independent experiments. C) GLI-dependent luciferase reporter assay. Shh-Light II cells were incubated for 22 hours with 30 nM SAG21k (SMO agonist) in the presence of different concentrations of **21**. The data are presented as mean ± SEM of three independent experiments. Cells in the absence of ligand had a mean value of −86.0 ±5.2 (data point not shown in the plot). D) Chemical structures of the prioritized hit compounds **2**, **6**, **21**, and **22** evaluated in the full concentration-range competition binding assays.

In summary, using the two experimental paradigms, we have confirmed that compound **21** *i*) binds to the core of the receptor and competes with BODIPY-cyclopamine, and *ii*) acts as a SMO antagonist blocking agonist-stimulated GLI signaling. In these assays, the compound was active at low micromolar concentrations.

### The identified SMO antagonists are selective c-MET inhibitors

For the 23 selected compounds, we proceeded with fixed-dose activity evaluation on c-MET kinase using 20 μM as a test concentration (Table 1). Among the tested compounds, **18**, **21**, and **33** inhibited the c-MET kinase activity by 98% or more at 20 μM. While these three compounds evoked a consistent inhibition of the kinase activity at a fixed dose concentration, only **21** was confirmed to antagonize SMO. Thus, **21** was selected for further IC_50_ determination using the pan-kinase modulator staurosporine as a reference for c-MET. As reported in Table 1, **21** showed an IC_50_ for c-MET in the sub-micromolar range (pIC_50_ 6.94 ± 0.07, reported as SEM).

**Table 1.** Biochemical evaluation of compounds **1-39**.

| Number | SMO pK<sub>i</sub> | c-MET % Inhibition<br>@20 µM | c-MET<br>pIC<sub>50</sub> |
| --- | --- | --- | --- |
| **1** | n.a. | 1.83 |  |
| **2** | 5.03 ± 0.43 | 8.50 |  |
| **3** | n.a. | 1.28 |  |
| **4** | n.a. | 3.61 |  |
| **5** | n.a. | 0.03 |  |
| **6** | 3.55 ± 0.97 | 14.70 |  |
| **10** | n.a. | 0.24 |  |
| **13** | n.a. | 5.78 |  |
| **14** | n.a. | 6.79 |  |
| **16** | n.a. | 6.90 |  |
| **17** | n.a. | 4.75 |  |
| **18** | n.a. | 100.42 |  |
| **20** | n.a. | 13.27 |  |
| **21** | 5.56 ± 0.21 | 99.10 | 6.94 ± 0.07 |
| **22** | 5.00 ± 0.08 | 19.80 |  |
| **23** | n.a. | 1.31 |  |
| **25** | n.a. | -0.87 |  |
| **26** | n.a. | 0.22 |  |
| **28** | n.a. | 10.73 |  |
| **31** | n.a. | 9.51 |  |
| **32** | n.a. | 5.29 |  |
| **33** | n.a. | 98.05 |  |
| **39** | n.a. | 14.41 |  |
| **SANT-1** | 7.93 ± 1.02 |  |  |
| **Staurosporine** |  | 96.3 | 7.29 ± 0.05 |

To explore the selectivity profile of **21**, the latter was also tested for its inhibition against a panel of 20 kinases, representative of the TKinome, including our main target c-MET. The results of this inspection are reported in Table 2. From this analysis, **21** emerges as the most proficient SMO antagonist/c-MET selective inhibitor with residual inhibitory activity on the receptor tyrosine kinases FGFR1 and TRKA.

**Table 2.** pIC_50_ evaluation of **21** against a panel of 20 TKs.

| Kinase | pIC<sub>50</sub> |  |
| --- | --- | --- |
|  | **21** | **Staurosporine** |
| ABL | > 4.70 | 7.11 ± 0.03 |
| CSK | > 4.70 | 7.49 ± 0.04 |
| EGFR | > 4.70 | 7.22 ± 0.08 |
| EPHA2 | > 4.70 | 7.00 ± 0.09 |
| EPHB4 | > 4.70 | 6.82 ± 0.06 |
| FGFR1 | 5.76 ± 0.02 | 8.69 ± 0.05 |
| FLT3 | > 4.70 | 9.91 ± 0.03 |
| IGF1R | > 4.70 | 7.56 ± 0.05 |
| ITK | > 4.70 | 8.61 ± 0.07 |
| JAK3 | > 4.70 | 9.74 ± 0.02 |
| KDR | 5.24 | 8.60 ± 0.04 |
| LCK | > 4.70 | 9.01 ± 0.08 |
| c-MET | 6.94 ± 0.07 | 7.29 ± 0.05 |
| PDGFRα | > 4.70 | 9.72 ± 0.02 |
| PYK2 | > 4.70 | 8.84 ± 0.05 |
| SRC | > 4.70 | 8.42 ± 0.05 |
| SYK | > 4.70 | 9.57 ± 0.07 |
| TIE2 | > 4.70 | 7.15 ± 0.04 |
| TRKA | 5.44 ± 0.04 | 9.36 ± 0.03 |
| TYRO3 | > 4.70 | 8.84 ± 0.09 |

### Docking analysis

To rationalize the promising dual activity of **21**, observed in the biological assays, in-depth structural analysis of its predicted binding modes was conducted. By carefully examining the docking poses within the binding sites of both c-MET and SMO, we sought to elucidate the key protein-ligand interactions driving its polypharmacological profile. Specifically, the analysis of the docking results for c-MET revealed that compound **21** effectively adopts an elongated conformation, corroborating its classification as a Type II inhibitor by PyRMD. This extended pose allows the quinoline nucleus to interact with the M1160 backbone amine via H-bond interactions (Fig. 3A and 3C). Additionally, the fluorophenyl ring is well positioned to form π-π interactions with the F1223 side chain. Furthermore, the amide of the benzamide forms H-bonds to the D1222 backbone carbonyl, while the methylpyrimidine might establish T-shaped interactions with the F1134 side chain. Concerning SMO, docking calculations revealed that the quinoline group establishes π-π interactions with the F484^6.65^ side chain (Fig. 3B and 3D). Moreover, the fluorophenyl ring, in the predicted binding mode, might form positive contacts with the adjacent N219 side chain, while the benzamide amine interacts via H-bond interactions with the D384 side chain located on the ECL2. Notably, the aminopyrimidine amine forms H-bonds with the Y394 side chain while its aromatic nucleus is well positioned in a receptor portion where favorable interactions with W281^2.58^, F391, R400^5.43^, and E518^7.38^ can be established.

**Fig. 3.**
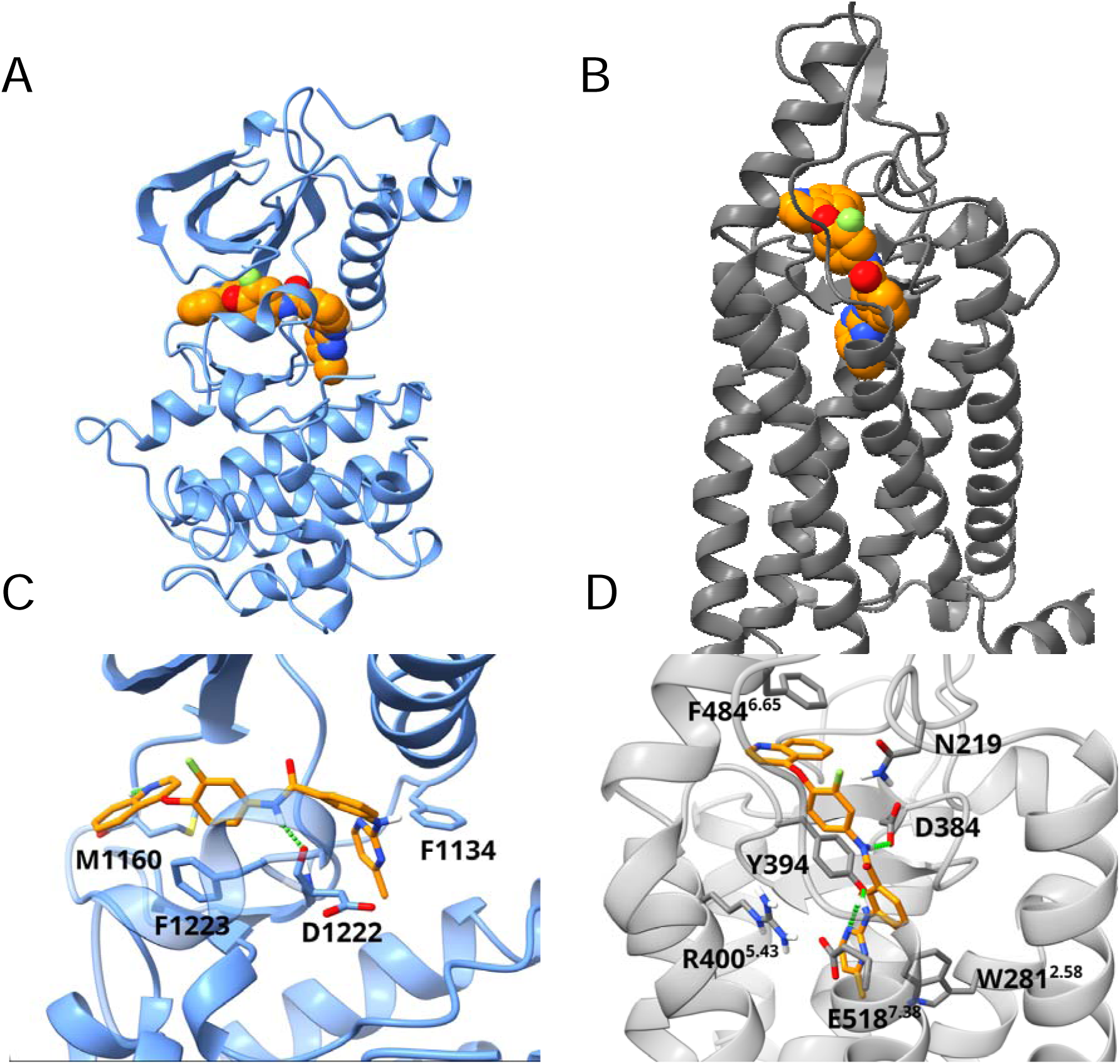
Predicted binding modes of compound 21 in complex with c-MET and SMO. (A, C) Global and close-up views of compound **21** (orange sticks) docked into the c-MET receptor (PDB ID: 4MXC, blue ribbons). (B, D) Global and close-up views of compound **21** (orange sticks) docked into the SMO receptor (PDB ID: 5L7I, grey ribbons).

### Compound 21 is antiproliferative in PDAC cells

For cell-based experiments, **21** was synthesized as described in the Supplementary Material. The antiproliferative effect of c-MET-SMO inhibition was evaluated in MIAPaCa2 and PANC-1, human pancreatic cancer cell lines, and in human pancreatic ductal cell line HPDE6c7, at 24h, 48h, and 72 hours, comparing the effect of **21** with the MET inhibitors PHA-665752 and Sitravatinib, and the SMO antagonist Sonidegib (Fig. 4A). The results showed a similar profile at 24h, with IC_50_ values increasing in the following order: PHA-665752, Sonidegib, Sitravatinib, and **21** (Fig. 4B). At 48 and 72 hours, the IC_50_ value for **21** was reduced compared to Sonidegib and Sitravatinib. In contrast, PHA-665752 showed different trends between the two models, displaying higher potency in PANC-1 than in MIAPaCa-2 cells. Overall, while PHA-665752 remained the most potent compound, **21** demonstrated a significant time-dependent increase in activity. **21** was moderately efficient on human ductal cells HPDE6c7 and exhibited higher IC_50_ values compared to the other inhibitors at up to 72 hours, indicating a reduced effect in non-tumor cells. Then, GR_50_ and GRmax metrics, representing the potency and efficacy of the treatments, were compared (Fig. 4C). **21** showed a weak effect in both models during the first 24h. Over time, it evolved towards a potent and efficacious effect in MIAPaCa2 cells during the following 48h, maintaining efficacy at 72h. In PANC-1, however, its potency slightly increased at 48h with low efficacy, before returning to a weaker effect at 72h. PHA-665752 showed a similar response in both models, with a weak effect at 24h that improved to high efficacy at 48h and 72h. Sitravatinib and Sonidegib showed similar efficacy in both models throughout the course of the experiment. HPDE6c7 cells showed measurable potency but consistently low maximal efficacy across all treatments and time points, indicating limited antiproliferative effects in normal cells. In contrast, pancreatic cancer cells exhibited greater sensitivity and a pronounced, time-dependent increase in efficacy, supporting a clear therapeutic window and the selective antitumor activity of compound **21**.

**Fig. 4.**
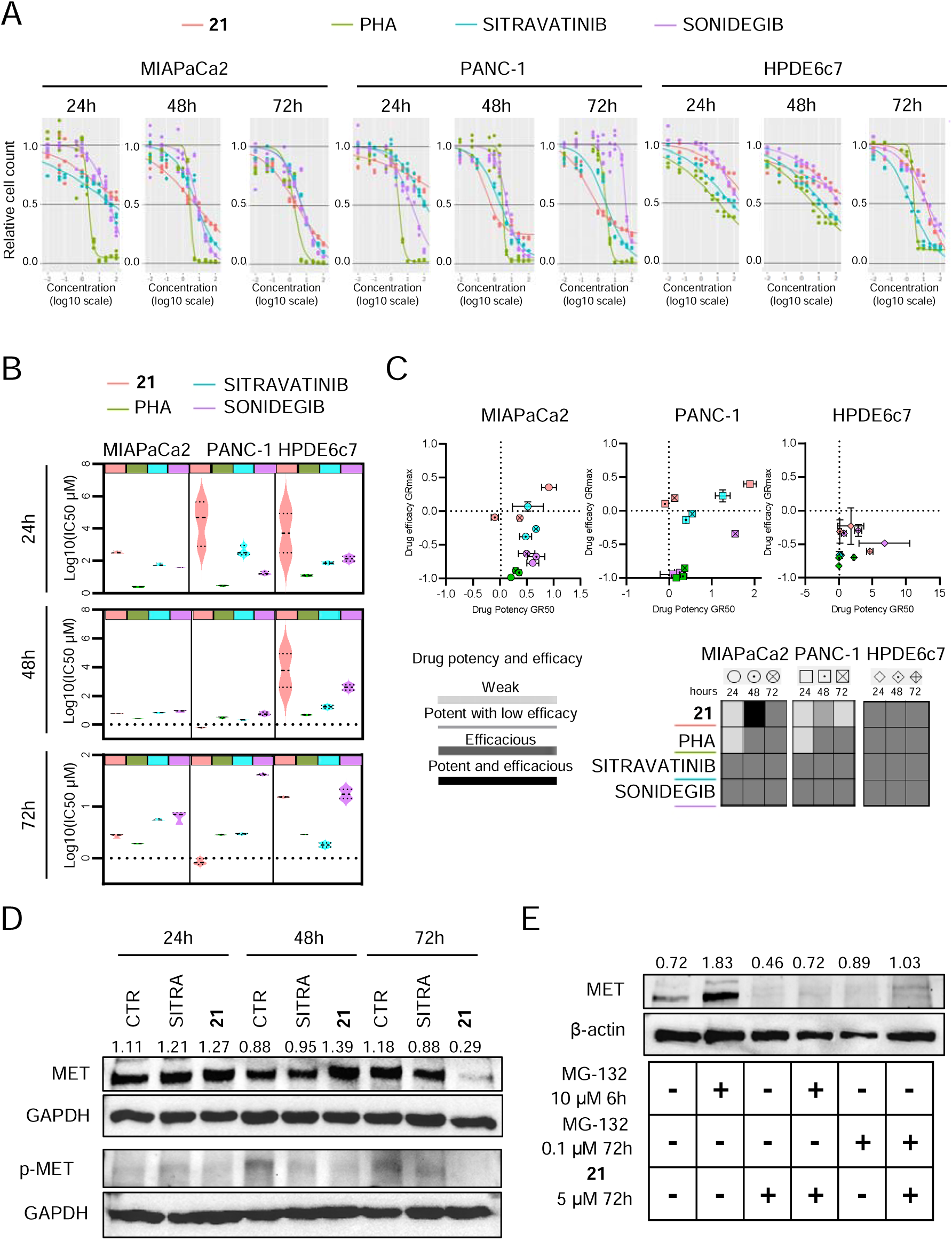
A) Viability assay in MIAPaCa2, PANC-1, and HPDE6c7 cells upon 21, PHA, Sitravatinib, and Sonidegib treatment in a time course of 24h, 48h, and 72h. X and y axis refer, respectively, to drug concentration in log10 and relative cell count normalized to control. B) Violin plots show IC_50_ in log10 value in MIAPaCa2, PANC-1, and HPDE6c7 upon **21**, PHA, Sitravatinib, and Sonidegib treatment at 24h, 48h, and 72h. C)Image indicates drug potency (GR50) in log10 and drug efficacy (GRmax) in MIAPaCa2 (circle),PANC-1 (square), HPDE6c7 (rhombus) upon 21, PHA, Sitravatinib, and Sonidegib treatment at 24h, 48h, and 72h. D) Western blot analysis shows MET and phosphorylated MET (p-MET) expression level normalized on GAPDH, in MIAPaCa2 exposed to Sitravatinib and 21, at 24h, 48h, and 72h. E) Western blot analysis shows MET expression level normalized on β-Actin, in MIAPaCa2 exposed to MG-132 at 10 µM for 6h and 0.1 µM for 72h, alone or in combination with **21** (5 μM) for 72h.

MET protein levels and phosphorylation were also investigated in MIAPaCa2 cells (Fig. 4D). Interestingly, upon treatment with **21**, total c-MET level drastically reduced at 72h, while the phosphorylated form progressively decreased starting from 48h. To investigate the mechanism underlying c-MET depletion, we assessed the role of the proteasome (Fig. 4E). Co-treatment with the proteasome inhibitor MG-132 restored c-MET protein levels, indicating that compound **21** promotes ubiquitin–proteasome-dependent degradation of the receptor. A similar effect has also been detected for Tivantinib, another c-MET inhibitor that facilitates its degradation (*33*). Building on this precedent, we hypothesize that the binding of compound **21** thermodynamically destabilizes the c-MET conformation, thereby promoting the chaperone-mediated recruitment of E3 ubiquitin ligases and subsequent proteasomal degradation.

### 21 blocks growth of 3D PDAC homo and heterospheres

To bridge the gap between two-dimensional cell cultures and the complex pathophysiology of pancreatic cancer, we utilized a customized 3D spheroid platform. Homotypic spheroids (MIAPaCa2) and heterotypic spheroids (incorporating CAF154-hTERT fibroblasts) were developed to test the hypothesis that **21** could overcome the fibroblast-mediated structural compaction and thus improve therapeutic efficacy. These studies were conducted by comparing the activity of **21** with the c-MET inhibitor PHA-665752 and the SMO antagonist Sonidegib. As shown in Fig. 5A, Sonidegib induced the appearance of a smooth, translucent peripheral hyaline halo, identified as cellular debris in MIAPaCa2 homotypic spheroids. PHA-665752 induced prominent surface roughness and debris shedding, indicating a toxic effect on peripheral cells. These effects were maximized by the combination of 100 µM Sonidegib + PHA-665752. In contrast, **21** triggered a massive release of debris at both 50 and 100 µM (Fig. 5A). Notably, the conventional projected cross-sectional area measurements failed to accurately reflect drug efficacy (Fig. 5B), as the loosening of the structure sometimes increased the diameter; however, efficacy was confirmed by a significant increase in LDH release (Fig. 5C).

**Fig. 5.**
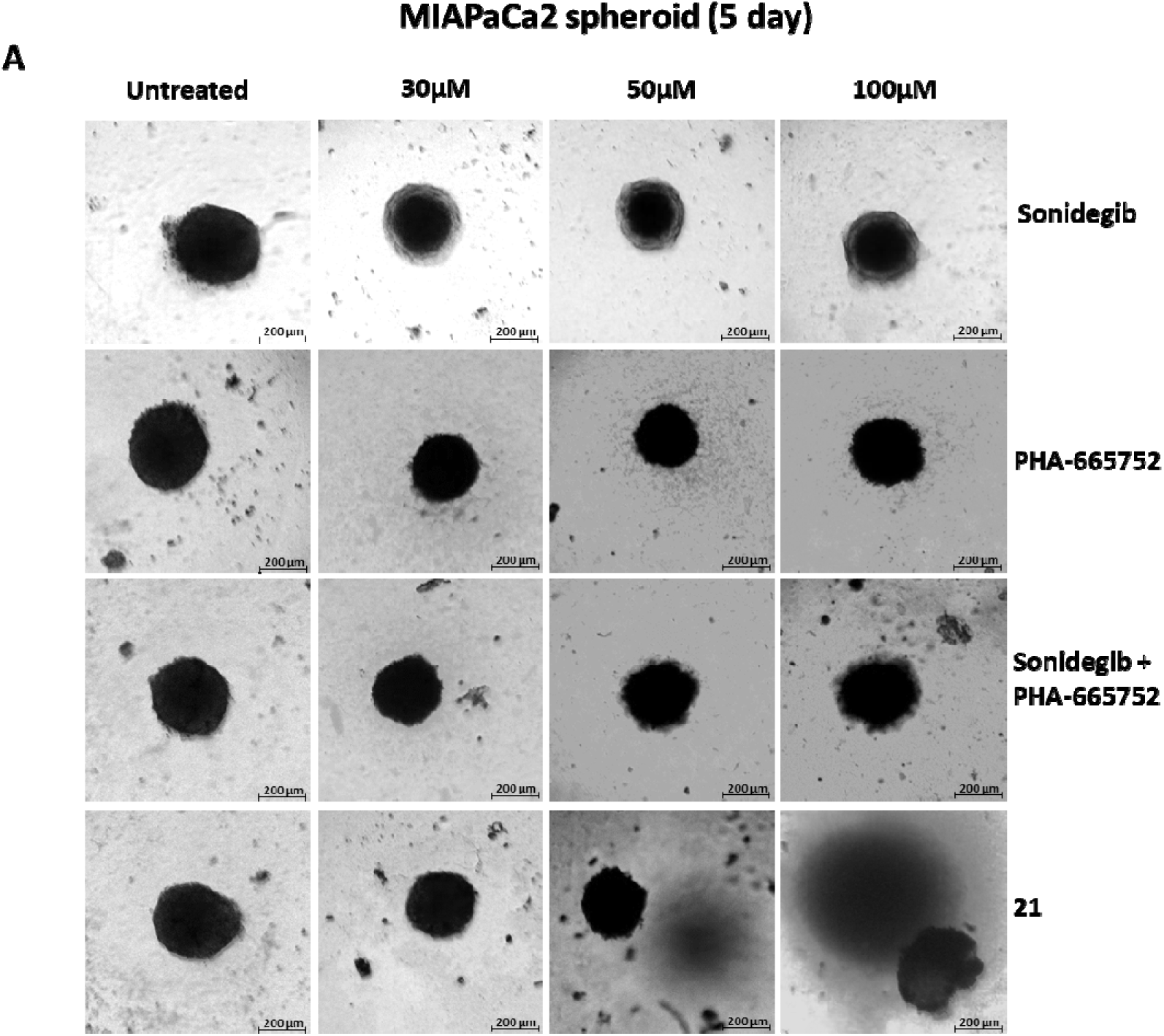
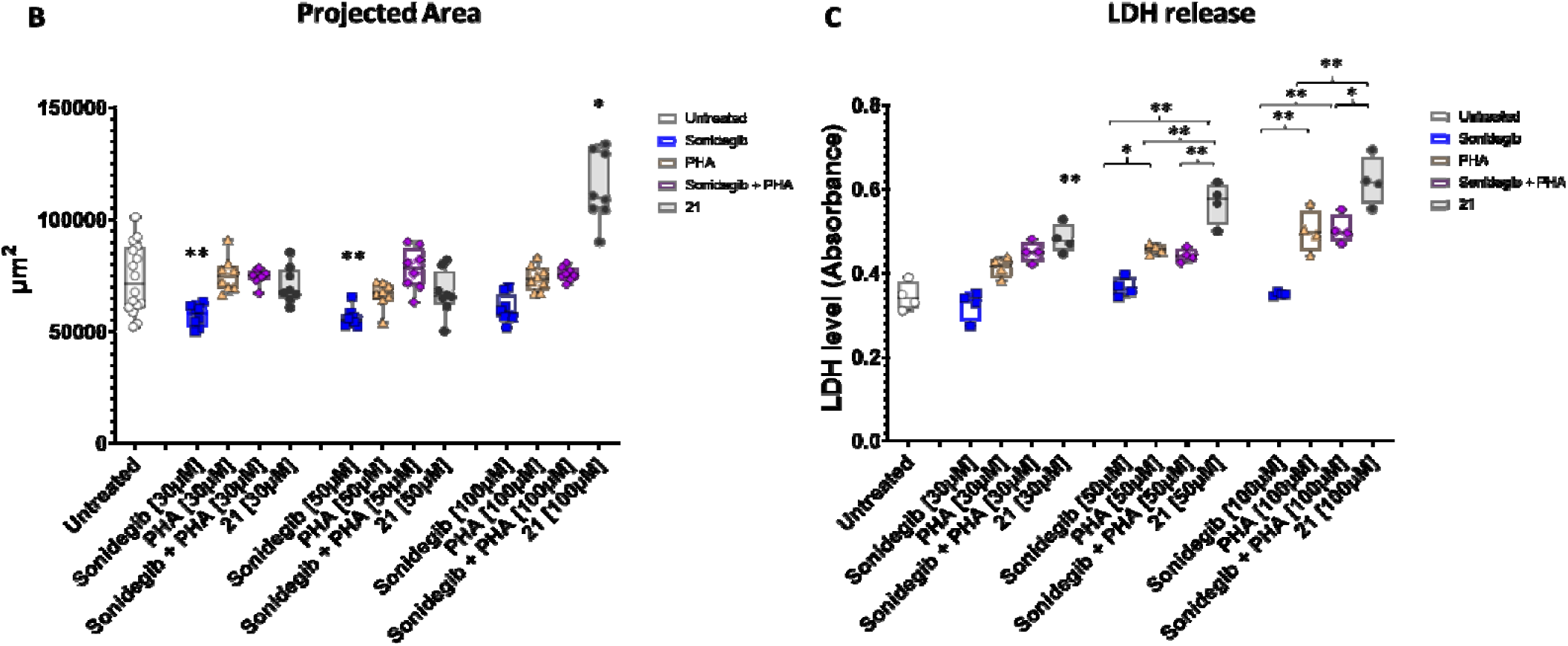
21 Exerts Potent Cytotoxic Effects on MIAPaCa2 homotypic 3D model. **(A)** Representative microscopy images (50×50 magnification) of MIAPaCa2 tumor spheroids at 120 hours (16 untreated control replicates and 8 treated replicates per condition). Scale bar: 200 µm. (**B)** Quantitative analysis of heterospheroid projected cross-sectional area (µm^2^) at 120 hours. **(C)** LDH release at 120 hours. Results were considered statistically significant at ANOVA with Šídák’s post-hoc test p < 0.05. All data are presented as mean ± standard deviation (SD).

In experiments shown in Fig. 6, the projected area of MIAPaCa2/CAF154-hTERT heterospheroid increased remarkably across all treated groups, counteracting the fibroblast-mediated compaction observed in untreated heterospheroids (Fig. 6A). Sonidegib exerted a localized effect on the outer 3D layers, resulting in volume expansion and the formation of a hyaline halo. PHA-665752-treated spheroids exhibited a loss of spherical edge and a more relaxed cellular architecture at 30 μM, with minimal debris. The Sonidegib + PHA-665752 combination induced dose-dependent surface roughness and an irregular perimeter, with increased debris (Fig. 6A). Crucially, in the **21**-treated group, massive, intense necrotic particulate debris surrounded the spheroids at all concentrations (30-100 μM) (Fig. 6A, lower panel).

**Fig. 6.**
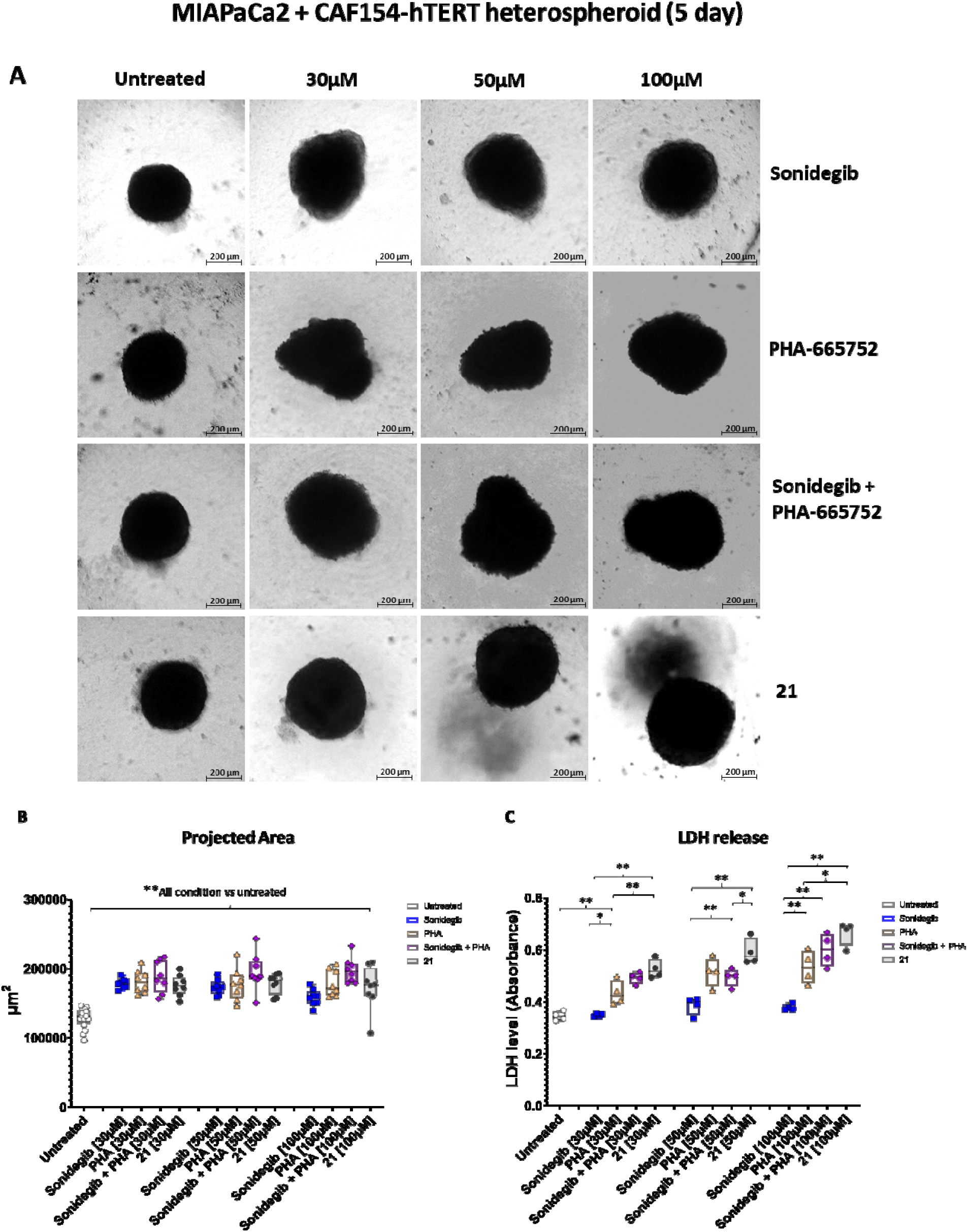
21 Exerts Potent Cytotoxic Effects on MIAPaCa2 + CAF154-hTERT 3D Models by overcoming the fibroblast-mediated structural cohesion. **(A)** Representative microscopy images (50x magnification) tumor MIAPaCa2 + CAF154-hTERT heterospheroid at 120 hours (16 untreated control replicates and 8 treated replicates per condition). Scale bar: 200 µm. **(B)** Quantitative analysis of heterospheroid projected cross-sectional area (µm^2^) at 120 hours. **(C)** LDH release at 120 hours. Results were considered statistically significant at ANOVA with Šídák’s post-hoc test p < 0.05. All data are presented as mean ± standard deviation (SD).

The structural loosening observed in treated heterospheroids was quantitatively confirmed by a substantial increase in projected area over time (128820.42±14481.29 µm^2^ in untreated vs 178869.20±6652.41 µm^2^ in 30µM Sonidegib, p=0.0001) (Fig. 6B). This loss of density correlated with higher LDH leakage (Fig. 6C). Across all tested concentrations, **21** consistently exhibited superior cytotoxicity compared to the Sonidegib + PHA-665752 combination (Fig. 6C).

To discriminate between epithelial and stromal cells within the heterospheroid structure, MIAPaCa2 and CAF154-hTERT cells were stained with fluorescent dyes. At 72 hours post-seeding, a highly organized spatial distribution was detectable (Fig. 7A). The heterospheroids displayed a characteristic “core-shell” architecture: a dense centralized cluster of MIAPaCa-2 cells (CellTrace™ Blue) surrounded by a distinct outer shell of CAF154-hTERT fibroblasts (CellTrace™ Far Red). As illustrated in Fig. 7B, compound treatments induced significant alterations to this architecture. While untreated hetero spheroids maintained an organized stromal shell after 8 days, Sonidegib caused only a moderate reduction in outer sheath compactness, leaving the tumor core relatively tight (compact blue signal as untreated). PHA-665752 began to affect the cancer cells, causing a “loosening” of the core (punctate blue signal). The combination showed only a slight additive effect. Remarkably, **21** elicited a severe disruption of the entire structure: both blue and red fluorescent signals appeared fragmented or “punctate”, with visible peripheral debris. This suggests that **21** acts by disrupting both fibroblast-fibroblast (CAF-CAF) and fibroblast-tumor interactions, implying a synergistic effect on spheroid integrity. These spatially balanced alterations are consistent with dual-compartment targeting, as supported by the morphological and toxicological data (Fig. 6C).

**Fig. 7.**
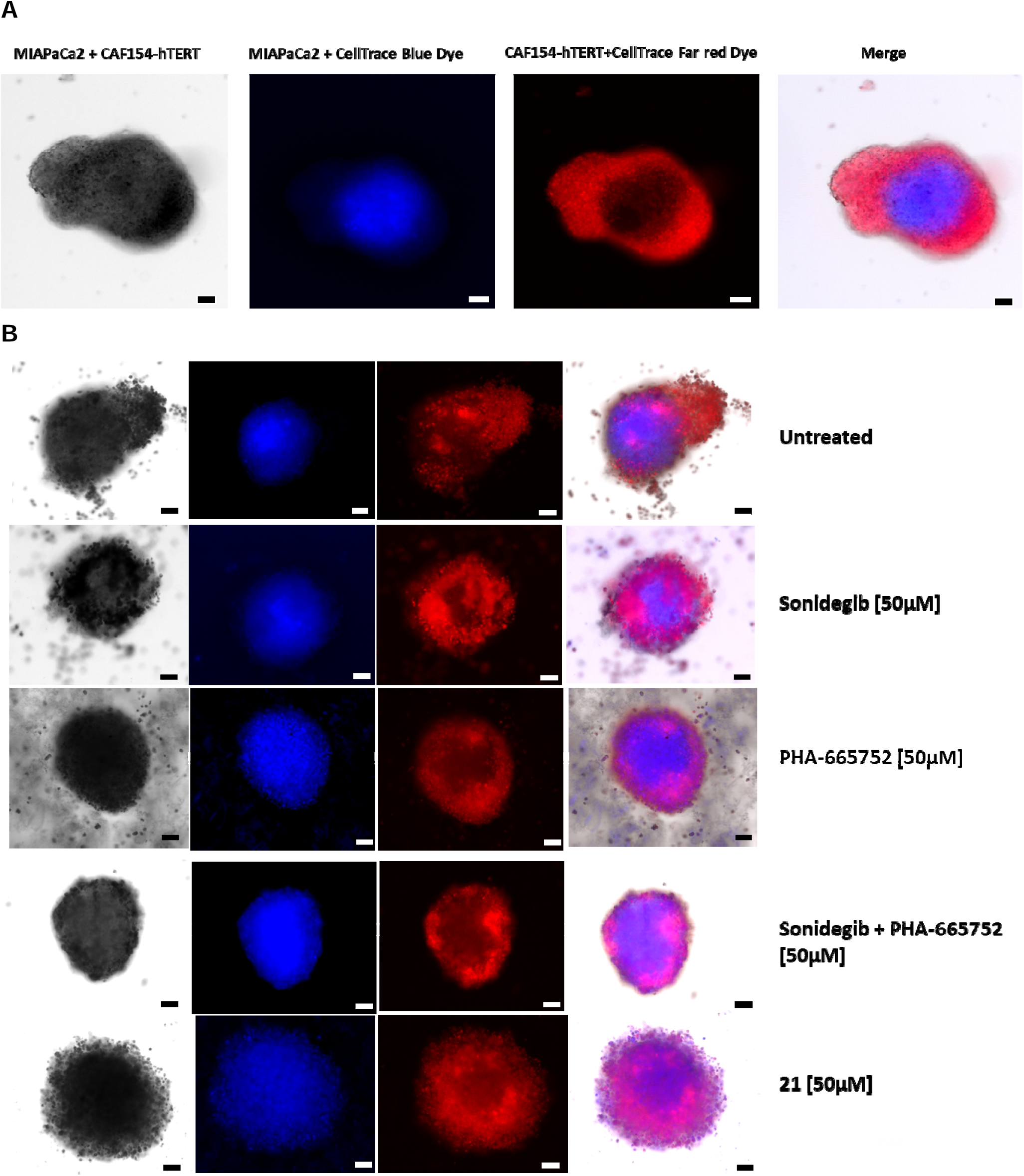
PDAC heterotypic 3D model showed Core-Shell Architecture and 21 treatment induced structural alteration. **(A)** Example of fluorescent labelled MIAPaCa2 and CAF154-hTERT 3D PDAC model (100x magnification Leica Mica Microhub widefield, microscopy). **(B)** Treatment-Induced Alterations in Heterospheroid Morphology (Day 8): Fluorescence microscopy analysis of heterospheroids after 72 hours of treatments. Black/white scalebar 50µm.

## DISCUSSION

The development of effective therapies for PDAC is severely hindered by the tumor’s complex microenvironment, which features dense desmoplasia and active paracrine signalling that drive profound drug resistance. In this setting, polypharmacology offers a promising strategy by simultaneously disrupting multiple interconnected oncogenic and stromal pathways with a single agent. In this study, we successfully employed an AI-guided virtual screening (VS) workflow to identify novel dual inhibitors of the c-MET and SMO-mediated pathways. Given that c-MET and SMO are phylogenetically and structurally distinct targets, accommodating the diverse chemical properties required to bind both pockets within a single molecule represents a significant design challenge.

To overcome this, we utilized PyRMD, a sophisticated ligand-based machine-learning tool developed by us. While the field of AI-driven drug discovery has increasingly shifted toward deep generative models and AlphaFold-driven de novo design, these platforms currently suffer from high rates of “hallucinated” molecules that are synthetically intractable or thermodynamically unstable. In contrast, our pipeline effectively filtered a highly curated, available library of approximately 9 million in-stock compounds down to 23 top candidates. This approach ensured rapid translation from in silico algorithmic prediction to in vitro biological validation, culminating in the identification of compound **21**, an aminopyrimidine-benzamide-phenoxyquinoline derivative that is obtained utilizing readily available starting materials and established synthetic methodologies.

Biochemical characterization confirmed that **21** acts as a highly active dual-modulator. It targets the orthosteric binding site of SMO, functionally antagonizing agonist-induced canonical signaling in the low micromolar range, while concurrently demonstrating potent, sub-micromolar inhibition of c-MET (pIC_50_ = 6.94 ± 0.07). Crucially, against a panel of 20 representative tyrosine kinases, **21** exhibited a highly favourable selectivity profile, sparing the vast majority of the tested TKinome. This selectivity is an essential characteristic for minimizing off-target toxicities in polypharmacological design.

The therapeutic potential of compound **21** was further supported in 2D cellular models of human PDAC. While initial anti-proliferative efficacy at 24 hours was weak, compound **21** exhibited a striking time-dependent increase in potency and efficacy at 48 and 72 hours. This sustained biological activity correlates strongly with its profound mechanism of action. In fact, Western blot analysis revealed that **21** drastically diminishes total c-MET protein levels over 72 hours. The complete rescue of c-MET expression upon co-treatment with the proteasome inhibitor MG-132 confirms that **21** triggers ubiquitin-proteasome-mediated degradation of the receptor.

Under physiological conditions, c-MET turnover is driven by ligand-induced autophosphorylation, which serves as the canonical signal to recruit E3 ubiquitin ligases (such as c-Cbl) for receptor internalization and degradation (*34*). Because compound **21** acts as a potent kinase inhibitor, it is expected to prevent autophosphorylation and thereby stabilize the inactive receptor at the cell membrane. However, our data indicate that compound **21** circumvents this standard physiological loop. We hypothesize that the specific extended binding mode of **21** within the ATP pocket induces a thermodynamic destabilization of the kinase domain. This structural disruption effectively mimics a misfolded state, forcing chaperone-mediated E3 ligase recruitment and subsequent proteasomal clearance despite the absence of canonical autophosphorylation.

Furthermore, bypassing the c-Cbl-dependent degradation pathway positions compound **21** as a highly promising candidate for targeting therapy-resistant MET mutants. Recent biochemical analyses highlight that the juxtamembrane domain of c-MET, specifically the region encoded by Exon 14, harbors the critical Y1003 docking site required for c-Cbl-mediated receptor turnover. In several malignancies, including pancreatic cancer (*35*), oncogenic Exon 14 skipping mutations physically eliminate this c-Cbl binding site, allowing the receptor to evade lysosomal degradation and drive prolonged, unchecked signaling (*36*). Because compound **21** induces degradation via structural destabilization and chaperone-mediated clearance, wholly independent of the Y1003/c-Cbl axis, it has the theoretical potential to effectively degrade and neutralize these highly aggressive, Exon 14-skipped MET variants that are recalcitrant to standard physiological turnover. Crucially, the therapeutic utility of inducing c-MET degradation relies entirely on kinase selectivity. While early-generation agents like tivantinib were once theorized to induce MET degradation, their clinical effects were ultimately attributed to non-specific, off-target cytotoxicity that occurred independent of sole c-MET engagement (*37*). In stark contrast, our kinome profiling establishes that compound **21** is a selective targeted agent. By specifically hijacking the ubiquitin-proteasome system, **21** functions analogously to modern targeted protein degradation (TPD) technologies, achieving complete target removal rather than only catalytic inhibition.

This target removal provides massive advantages over traditional competitive inhibitors such as PHA-665752. First, maintaining inactive c-MET at the membrane allows tumor cells to orchestrate bypass resistance by providing a physical docking station for HER3 transactivation (*38*) and AXL heterodimerization (*39*). Furthermore, intact c-MET physically tethers CD44v6 and integrin complexes to drive metastatic plasticity(*39*). By physically degrading the c-MET protein, compound **21** should dismantle these critical cytoarchitectural hubs, precluding the most aggressive avenues of acquired resistance. Second, degrading c-MET severs a newly discovered, immune-independent oncogenic dependency in PDAC, the endogenous PD-1/c-MET axis (*17*). By eliminating c-MET, compound **21** should neutralize this tumor-intrinsic signaling loop, offering a valuable strategy for overcoming resistance in patients who fail standard immune checkpoint inhibitors.

This unique targeted degradation mechanism complements the simultaneous SMO modulation required to tackle the desmoplastic stroma. High-resolution transcriptomics reveal that the PDAC stroma is highly heterogeneous; while SMO inhibition successfully depletes tumor-restraining myofibroblastic CAFs (myCAFs) to alleviate physical barriers (*40*), it paradoxically triggers a compensatory shift that enriches inflammatory CAFs (iCAFs) (*22*). These iCAFs secrete a pro-inflammatory cytokine milieu that activates the HGF/c-MET axis on tumor cells, driving rapid epithelial-to-mesenchymal transition (EMT) and accelerated metastasis, a phenomenon that has historically caused single-agent SMO inhibitors to fail in the clinic. Because compound **21** concurrently eradicates c-MET, it acts as a molecular safeguard, effectively neutralizing the tumor cells’ ability to respond to iCAF-mediated inflammatory surge.

To validate this synchronized blockade clinically, we employed a 3D heterotypic PDAC model comprising MIAPaCa2 cells and an outer shell of CAF154-hTERT fibroblasts. In these mature heterospheroids, compound **21** achieved superior cytotoxicity compared to the combination of individual inhibitors (Sonidegib + PHA-665752). We observed a lack of significant dimensional shrinkage following treatment due to the high structural resilience of the fibroblast-secreted extracellular matrix (ECM). The ECM maintains external volume even as internal cancer cells undergo massive necrosis; consequently, morphological alterations (progressive decompaction and debris shedding) alongside LDH release proved to be highly sensitive indicators of internal tumor disintegration.

Our core-shell spheroids replicate the physical desmoplastic barrier and localized paracrine crosstalk (*41, 42*), where the outer layer of CAFs replicates the dense desmoplastic stroma characteristic of human PDAC, serving as a physical barrier that limits drug penetration. Furthermore, the close contact between the inner MIAPaCa2 core and the outer CAF154-hTERT shell enables critical paracrine signaling and tumor-stroma crosstalk, driving the activation of multiple resistance pathways (*43*). Nevertheless, this ex vivo system lacks systemic immune and vascular compartments. Future validation of compound **21** should leverage advanced microfluidic “tumor-on-a-chip” platforms that incorporate endothelial microchannels and patient-derived immune populations. Such models will be crucial to fully quantify **21**’s ability to alleviate interstitial fluid pressure, reverse T-cell exclusion, and ultimately translate this dual SMO/c-MET inhibition strategy into a highly effective intervention for pancreatic cancer. Furthermore, while standard in vivo cell-line-derived xenograft (CDX) and patient-derived xenograft (PDX) models are frequently used to evaluate novel therapeutics, they present a profound limitation for studying tumor-stroma polypharmacology. It is well established that upon engraftment in immunodeficient mice, human stromal populations are rapidly outcompeted and entirely replaced by murine host stroma (*44, 45*). This murine stromal replacement fundamentally disrupts the species-specific paracrine signaling loop required to evaluate our mechanism of action. Specifically, it creates a severe species mismatch wherein human tumor-derived Hedgehog ligands must activate murine SMO on host CAFs, which in turn must secrete murine cytokines to activate human c-MET on the neoplastic cells. Because compound **21** was specifically designed to target the human SMO and human c-MET binding pockets, testing it in a chimeric human/murine environment would yield inherently confounded results due to cross-species receptor incompatibilities. Therefore, the completely humanized 3D heterotypic core-shell spheroids utilized in this study (MIAPaCa2 and CAF154-hTERT) do not merely serve as an in vitro proxy; they represent a strictly necessary, superior physiological model to preserve and accurately evaluate the intact human-human tumor-stroma crosstalk.

## CONCLUSION

In summary, this study validates the utility of an AI-driven virtual screening workflow in identifying compound **21**, an aminopyrimidine-benzamide-phenoxyquinoline derivative, as a novel, selective dual modulator of the SMO and c-MET pathways. Beyond its potent dual affinity, **21** distinguishes itself from traditional competitive inhibitors by inducing the proteasomal degradation of c-MET. This crucial mechanism entirely removes the structural scaffold required for non-canonical bypass signaling, thereby conferring a prolonged therapeutic window. Furthermore, our validation in complex 3D heterotypic models demonstrated that this single-molecule polypharmacological approach yields superior cytotoxicity compared to the combination of selective single-target inhibitors. By synchronizing the blockade of SMO-mediated stromal support and MET-driven invasive potential, compound **21** effectively overcomes the structural resilience of the desmoplastic stroma while circumventing the risks of discordant drug distribution and paradoxical EMT induction often associated with combination therapies. Ultimately, these findings establish dual SMO/c-MET inhibition as a robust, highly translational therapeutic strategy for tackling the intractable microenvironment of pancreatic ductal adenocarcinoma.

## MATERIALS AND METHODS

### AI-enforced VS

To create a machine learning (ML) prediction model, PyRMD was fed with a comma-separated file (.csv) retrieved from the ChEMBL database for c-MET. This led to the construction and preparation of the training dataset. According to their bioactivity data, the compounds included in the .csv file were classified into three groups: actives, inactives, and discarded. For benchmarking purposes, several activity thresholds (i.e., 101–1001 nM for the “actives”) were systematically evaluated. Specifically, compounds whose bioactivity fell below the selected activity threshold were placed in the “actives” group. In contrast, those with reported bioactivity exceeding the inactivity threshold of 40,000 nM were assigned to the “inactives” group. By selecting the MinHash fingerprints (MHFP) for the featurization process and by varying the ε cutoff for “actives” and “inactives” (0.01-0.99 with a 0.01 step), 88209 different models were generated. For all models, PyRMD returns relevant metrics to evaluate their predictive performance (i.e., TPR, FPR, F-score, ROC AUC, BED ROC, PRC AUC). In this work, the selected model was chosen by maximizing the F-score value. Once the model was generated, PyRMD was used to screen the Mcule in-stock medium-large database (∼9 million compounds), and it automatically returns all the compounds deemed to be active along with a confidence score of its prediction (RMD Score)

### Molecular docking

For docking calculations, from the Protein Data Bank repository, two c-MET (type I and type II) and six SMO crystal structures were downloaded (PDB: 4R1V, PDB: 4MXC, and PDB: 4JKV, 4N4W, 4O9R, 4QIM, 5L7I, 5V57, respectively). All the structures were prepared for the docking calculations through the protein preparation wizard in Maestro. For each of them, the hydrogen atoms were added and minimized, the solvent molecules were removed, and the appropriate protonation and tautomeric state of the protein side chains were calculated. These structures were used for the docking calculations employing AutoDock Vina. As demonstrated in our recent retrospective analysis, applying AutoDock Vina across the ensemble of all the SMO receptors yielded the highest enrichment factors (EF) within the top 25% of the ranked database (*46*). Using the AutoDockTools Python scripts, the structures were converted to the AutoDock PDBQT format, where, compared to a standard PDB file, Gasteiger charges are added to the atoms, and the torsional freedoms of the various bonds are described. The docking grid box was centered on the binding site and made large enough to include the volume occupied by the co-crystal ligands. Then, the receptor grid maps were calculated with the AutoGrid4 software, mapping the receptor interaction energies using every AutoDock atom type as a probe. Each compound was docked one at a time in the protein structures employing standard settings. Once the docking calculations were completed, the docked lowest-energy poses and the largest cluster poses (the pose found most frequently in the docking experiment) were extracted as PDB files along with their predicted binding free energy.

### SMO binding and functional assays

ΔCRD Nluc-SMO cells (*47*) were seeded onto a flat-bottom black 96-well plate (Greiner or ThermoFisher). 24 hours later, the cells were washed once with 200 μl of HBSS (HyClone) and incubated for 30 minutes at 37 °C with CO_2_ in 70 μL of HBSS with the compounds, followed by the addition of 10 μL of BODIPY-cyclopamine (Biosynth) at a final concentration of 5 nM (the screening experiments) or 10 nM (the experiments with a full concentration range). Subsequently, the cells were incubated at 37 °C with CO_2_ for 90 min. Finally, 10 μL of 1:1000 final dilution of furimazine (Promega) was added, and the plate was incubated at 37 °C without CO_2_ for 10 min. The raw BRET ratio is defined as the ratio of BRET acceptor (BODIPY; 520–560 nm) emission signal over the BRET donor (Nluc; 460–500 nm) emission signal. %ΔBRET ratio for every ligand concentration is calculated as the percentage of difference in the BRET ratio of cells treated with the ligands together with BODIPY-cyclopamine and BODIPY-cyclopamine-treated cells, divided by the BRET ratio of BODIPY-cyclopamine-treated cells. The BRET measurements were recorded with a 100 ms integration time using a TECAN Spark microplate reader (TECAN). A high-potency SMO antagonist, SANT-1, was used as a positive control (Abcam).

Inhibition of SMO-mediated signaling was assessed using the GLI Luciferase Reporter Shh Light II cell line, a NIH/3T3 cell line-derived HH reporter cell line that stably expresses a GLI-responsive Firefly luciferase reporter (Fluc) and constitutive *Renilla* luciferase (*R*luc) to measure HH pathway activation (a kind gift from Rune Toftgård). The cells were seeded onto white 96-well plates, and following a 6-hour incubation, different concentrations of the ligand **21** together with 30 nM of the SMO agonist SAG21k (Abcam) were added. The cells were incubated for a further 22 hours at 37 °C with CO_2_. Bioluminescence was measured using the Dual-Luciferase Reporter Assay System (Promega) with a TECAN Spark microplate reader. Cells treated with SAG21k at 30 nM only were used as a positive control (baseline). Data are presented as % the difference (Δ) of Fluc/*R*luc luminescence ratio = *100%*(ligand 21-SAG21k alone)/SAG21k alone* for each ligand 21 concentration.

### c-MET inhibition assays and selectivity profile

For fixed-dose inhibition of ABL, CSK, EGFR, EPHA2, EPHB4, FGFR1, FLT3, IGF1R, ITK, JAK3, KDR, LCK, c-MET, PDGFRα, PYK2, SRC, SYK, TIE2, TRKA, and TYRO3 TKs and IC_50_ c-MET inhibition studies, the test compounds were dissolved in DMSO to achieve a 2 mM concentration. Then the solution was further diluted with assay buffer to make the final test compound solutions. Reference compounds for assay control were prepared similarly. An off-chip mobility shift assay (MSA) was then used to test the inhibition of the compounds against the selected TKs. Briefly, the 4x substrate/ATP/metal solution was prepared with kit buffer (20 mM HEPES, 0.01% Triton X-100, 5 mM DTT, pH 7.5), and the 2x kinase solution was prepared with assay buffer (20 mM HEPES, 0.01% Triton X-100, 5 mM DTT, pH 7.5). 5 μL of 4x compound solution, 5 μL of 4x substrate/ATP/metal solution, and 10 μL of 2x kinase solution were mixed and incubated in a well of a polypropylene 384-well microplate for 1h at RT. 70 μL of termination buffer (QuickScout Screening Assist MSA; Carna Biosciences) was added to the well. The reaction mixture was applied to the LabChipTM system (Perkin Elmer), and the product and substrate peptide peaks were separated and quantified. The kinase reaction was evaluated by the product ratio calculated from the peak heights of product (P) and substrate (S) peptides (P/(P+S)). The readout value of reaction control (complete reaction mixture) was set to 0% inhibition, and the readout value of background (enzyme(-)) was set as 100% inhibition; then, the percent inhibition of each test solution was calculated. IC_50_ value was calculated from concentration vs %Inhibition curves by fitting to a four-parameter logistic curve. All experiments were run in triplicate and results expressed as averages ± SD.

### Cell-based experiments

*Cell Lines.* MIAPaCa2 (CRL-1420) and PANC1 (CRL-1469) cell lines were purchased from ATCC (Milan, Italy). Cell lines were grown in DMEM (Euroclone, #ECB7501L), supplemented with 10% heat-inactivated fetal bovine serum (Sigma-Aldrich, #F7524), penicillin–streptomycin mix respectively used at 100 U/mL and 100 μg/mL (Euroclone, #ECB3001D), 250 ng/mL amphotericin B (Euroclone, ECM0009D), and 2 mM l-glutamine (Euroclone, ECB3000D). HPDE6c7 cells were maintained in Advanced DMEM F12 (Gibco, #12634-010), B27 17504044 (Gibco, #17504044), N2 (Gibco, #17502-048), bFGF (PEPROTECH, #AF-100-18b-250UG), EGF (PEPROTECH, #AF-100-15-500UG), Hepes (Euroclone, # ECM0180D). Glutamax (Gibco, #35050-038). Cells were cultured with incubator parameters: 37 °C with 5% CO2. Cells were analyzed for mycoplasma contamination with EZ-PCR Mycoplasma Test Kit (Biological Industries, #20–700-20). We performed a heterotypic 3D screening employing the non-commercially available, patient-derived cancer-associated fibroblast (CAF) cell line, CAF154-hTERT (*48*). CAF154-hTERT cells were cultured in Fibroblast Basal Medium (PCS-201-030^TM^) supplemented with Fibroblast Growth Kit–Low Serum (PCS-201-041^TM^) to ensure a standardized culture environment.

### Cell viability assay

Cell viability assay in MIAPaCa2, PANC-1, and HPDE6c7 cell lines was assessed using thiazolyl blue tetrazolium bromide (MTT; Sigma-Aldrich, #57,360–69-7) according to the manufacturer’s instructions. 3[×[10^3^ cells/well were seeded in a 96-well plate. The following day, cells were treated with 21, PHA-665752 (MedChemExpress, #HY-11107), Sitravatinib (Fisher Scientific, #T4349), and Sonidegib (Selleck Chemicals, #NVP-LDE225) used at final concentrations of 0.01-0.05-0.5-1-2-5-10-25-50-100 µM for 24h, 48h, and 72h. Absorbance values were measured at a wavelength of 570 nm using Infinite M-plex (Tecan, 30,190,085). Calculation of drug potency and efficacy, respectively indicated as GR50 and GRmax, and IC50 was conducted using GRmetrics (PMID: 29065900) as an R program with RStudio version 4.4.1. Interpolation of GR50 and GRmax values was used to assign effect: weak, potent with low efficacy, efficacious, and potent and efficacious.

### Western Blot analysis

MIAPaCa2 cells have been treated with **21** at 5 μΜ to investigate MET and ph-MET modulation using Sitravatinib at 10 μΜ as MET-inhibitor reference compound, in a time course of 24h, 48h, and 72h. To evaluate MET proteasomal degradation induced by 21 at 5 μΜ for 72h, MIAPaCa2 cells were treated with MG-132 (sc-201270, Santa Cruz Biotechnology) at 10 μΜ for 6h and 0.1 μΜ for 72h in combination and alone. Following, cell pellet extraction was performed as described before (*49*). Then, 50 μg of cell extract was loaded on 10% polyacrylamide gel for electrophoretic separation and transferred onto nitrocellulose membranes. Primary antibodies: MET (D1C2) mAb #8198, Phospho-MET (Tyr1234/1235) (D26) mAb #3077, GAPDH (14C10) Rabbit mAb #2118 were purchased from Cell Signaling Technology, and β-Actin (ACTBD11B7) from Santa Cruz Biotechnology. Antibodies were used following the datasheet protocol. Immunoreactivity was detected with Horseradish peroxidase-conjugated secondary antibodies (Bio-Rad, anti-rabbit, #1,705,046, anti-mouse, #1,706,516), and chemiluminescence was detected using the ECL reagent (Clarity Western ECL Substrate, 500 ml #1,705,061). Chemiluminescent signals were acquired using the Bio-Rad ChemiDoc Imaging System, and band intensities were quantified using ImageJ software (version 1.44).

### Imaging

GraphPad Prism version 8.3.0 was used for data visualization.

### Pancreatic Homotypic and Heterotypic Spheroids Development

Pancreatic heterotypic 3D cultures consisted of MIAPaCa2 human pancreatic tumor cells and cancer-associated fibroblast CAF154-hTERT. MIAPaCa2 cells were seeded in 96-well round-bottom ultra-low attachment (ULA) plates at a density of 3 × 10^3 cells/ well and incubated at 37 °C. After 48 hours, CAF154-hTERT cells were added to the MIAPaCa2 spheroids at a density of 9 × 10^3 cells/ well (*43*). Following an additional 24 hours, MIAPaCa2 spheroids or MIAPaCa2 + CAF154-hTERT heterospheroids were treated with Sonidegib, PHA-665752, their combination, or compound **21** at concentrations of 30 µM, 50 µM, or 100 µM. Digital micrographs were captured at 24, 96, and 120 hours using an inverted optical microscope (10x objective) (Axiovert 100, Carl Zeiss, Germany). ImageJ software was employed to quantify the projected cross-sectional area of each 3D culture.

### LDH Cytotoxic Assay

The release of Lactate Dehydrogenase (LDH) into the supernatant after 120h of treatment was used as a marker of drug-induced cell cytotoxicity. The assay was performed using the CyQUANT LDH Cytotoxicity Assay (Invitrogen, Waltham, MA, USA) according to the manufacturer’s instructions. Absorbance (Optical density, OD) was measured using a Mithras LB 940 Multimode Plate Reader (Berthold Technologies).

### Spatial Tracking via Fluorescence Microscopy

For spatial tracking within the 3D model, MIAPaCa-2 and CAF154-hTERT cells were labeled with CellTrace™ Blue and CellTrace™ Far Red dyes (Invitrogen™), respectively, prior to sequential seeding. The labeling was performed according to the manufacturer’s instructions. Drug treatments (50 µM) were initiated after the addition of fibroblasts. The spatial distribution and growth kinetics of the 3D PDAC model were monitored via fluorescence microscopy using a Leica Mica Microhub (confocal and widefield modes) at 10x magnification.

### Statistical analysis

The competition binding data and the GLI reporter inhibition data were fitted to a three-parameter model using GraphPad PRISM 8. The error bars on the plots representing competition binding and the inhibition of GLI signaling represent mean ± SEM, whereas the error bars on the plots with the binding data from the screening experiments represent mean ± SD. Two independent competition binding screening experiments, three independent full-concentration range competition binding experiments, and three independent GLI signaling inhibition experiments were performed. Each independent experiment was performed in technical triplicate.

In Pancreatic heterotypic 3D assays, the statistical design consisted of 16 untreated control replicates and 8 treated replicates per condition to account for the inherent biological variability of the models. Data normality was assessed using the Shapiro-Wilk and Kolmogorov-Smirnov tests, confirming a normal distribution for all variables. Homogeneity of variances was verified by Levene’s test. Comparison between treatments was performed using ANOVA followed by Šídák’s post-hoc test. Results were considered statistically significant at p < 0.05, and all data are presented as mean ± standard deviation (SD).

## List of Supplementary Materials

Materials and Methods for the synthesis of compound **21**. Fig S1 to S3 for multiple supplementary figures

## Acknowledgments

The authors wish to thank Dr. Luca Roz (Tumor Genomics Unit, Milan, Italy) for providing the patient-derived, non-commercially available cancer-associated fibroblast (CAF) line CAF154-hTERT.

## Author contributions

### Conceptualization

GA DP RB PK SC; Methodology: SC; Software: MR; Validation: UC; Formal analysis: MR; Investigation: MR UC VA CV PK; Supervision: CI CD SDM FC FM SS LA GS RB; Funding acquisition: GA PK GS LA RB; Project administration: SC.

## Data and material availability

All data associated with this study are present in the paper or in the Supplementary Materials.

