## Supplementary Material for "Targeting the Tumor-Stroma Crosstalk: An AI-Based Virtual Screening Strategy for Dual MET/SMO Inhibitors in Pancreatic Cancer"

**^1^DiSTABiF, University of Campania Luigi Vanvitelli, 81100 Caserta, Italy**

**^2^Department of Precision Medicine, University of Campania "Luigi Vanvitelli", 80138 Naples, Italy.**

**^3^Department of Chemical, Pharmaceutical and Agricultural Sciences, University of Ferrara, 44121 Ferrara, Italy.**

**^4^Microenvironment Molecular Targets Unit, Istituto Nazionale Tumori - IRCCS -Fondazione G. Pascale, Napoli, Italy.**

**^5^Karolinska Institutet, Department Physiology & Pharmacology, Sec. Receptor Biology & Signaling, Biomedicum, Stockholm, Sweden.**

**Synthesis of 21.** Compound **21** was obtained as shown in Scheme 1 to explore its synthetic accessibility. Briefly, 2-chloro-4-methylpyrimidine was reacted with 3-aminobenzoic acid to yield the nucleophilic aromatic substitution product **21a**. The subsequent HATU-mediated coupling with **21c** led to the target compound **21**. Compound **21c** was synthesized starting from 2-fluoro-4-nitrophenol, which was treated with 4-chloroquinoline under reflux conditions to generate the diarylether intermediate **21b**, followed by catalytic hydrogenation to reduce the nitro group. Overall, the synthesis of **21** benefits from the use of readily available starting materials and established methodologies, indicating its amenability for facile preparation (Fig. S1).

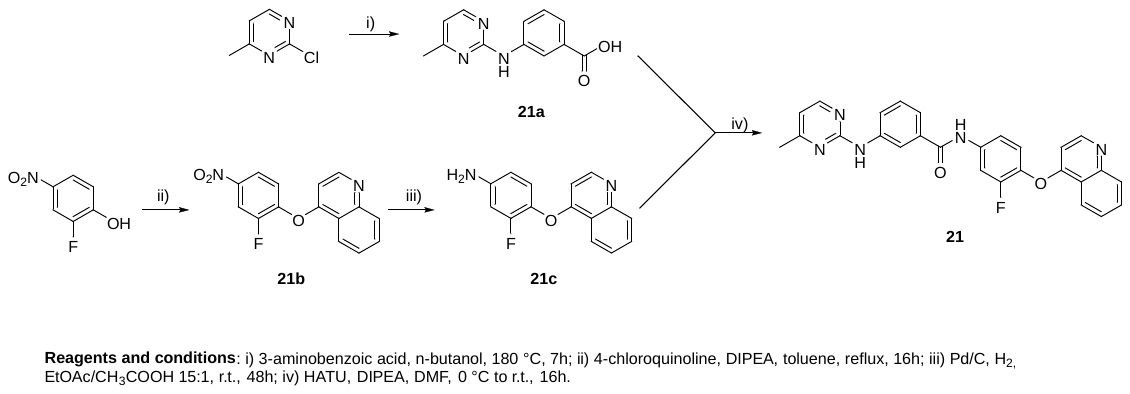

**Fig. S1**. Synthetic pathway for compound 21.

All reagents and solvents were sourced from BLD and Fluorochem. Reaction progress was monitored via thin-layer chromatography (TLC) using pre-coated Macherey-Nagel silica gel plates (F254, visualized under 254 nm UV light) and/or electrospray ionization mass spectrometry (ESI-MS) on a MICROMASS ZMD 2000. Analytical high-performance liquid chromatography (HPLC) was performed using a Beckman 116 liquid chromatograph coupled with a Beckman 166 diode array detector. Compound purity was assessed on a Phenomenex Kinetex C18 column (150 × 4.6 mm, 5 µm particle size) with a UV detector set to 220 nm. Compound **21** demonstrated > 95% purity by HPLC analysis (see HPLC analytical profile, Fig. S2). The mobile phase consisted of H₂O and CH₃CN, both containing 0.1% v/v trifluoroacetic acid (TFA), at a flow rate of 0.5 mL/min, employing a linear gradient from 0 to 100% CH₃CN over 25 minutes. Nuclear magnetic resonance (NMR) spectra (^1^H and ^13^C) were acquired on a Varian 400 MHz spectrometer. Chemical shifts (δ) are reported in parts per million (ppm), referenced to the residual ^1^H signal of the deuterated solvent (CDCl_3_: δ 7.26 ppm; DMSO-*d_6_*: δ 2.50 ppm). Coupling constants (*J*) are given in Hertz (Hz). Signal multiplicities are abbreviated as follows: s (singlet), d (doublet), t (triplet), q (quartet), m (multiplet), and b (broad). High-resolution mass spectrometry (HRMS) was performed using a Thermo Fisher UltiMate 3000 UHPLC coupled with an Orbitrap Q-Exactive HRMS.

*Synthesis of 3-((4-methylpyrimidin-2-yl)amino)benzoic acid (****21a****).* To a solution of 2-chloro-4-methylpyrimidine (1.94 mmol, 1.0 eq) in n-butanol, 3-aminobenzoic acid (1.94 mmol, 1.0 eq) was added at room temperature. The mixture was heated to 180 °C for 1 hour, after which the reaction progress was monitored via TLC (ethyl acetate/petroleum ether 1:3) and ESI-MS, both confirming the completion of the reaction. The crude mixture was then diluted with 1M NaOH (3 eq) and stirred for further 6 hours at room temperature. Subsequently, *n*-butanol was removed under vacuum, and the residue was acidified with 1M HCl (to pH 3-4), followed by extraction using ethyl acetate (3x10 mL). After solvent evaporation, a yellow solid was obtained (87% yield) and utilized in the next step without further purification.

^1^H NMR (400 MHz, DMSO-*d_6_*): *δ* 12.81 (bs, 1H), 9.72 (s, 1H), 8.45 – 8.28 (m, 2H), 8.00 (d, *J* = 8.1 Hz, 1H), 7.49 (d, *J* = 7.5 Hz, 1H), 7.36 (t, *J* = 7.9 Hz, 1H), 6.75 (d, *J* = 5.0 Hz, 1H), 2.35 (s, 3H). MS (ESI): m/z calcd for C_12_H_11_N_3_O_2_ [M + H]^+^ 230.25; found 230.29.

*Synthesis of 4-(2-fluoro-4-nitrophenoxy)quinoline (****21b****).* To a solution of 4-chloroquinoline (6.11 mmol, 1.0 eq) in toluene, 2-fluoro-4-nitrophenol (9.17 mmol, 1.5 eq), and DIPEA (18.34 mmol, 3.0 eq) were added. The mixture was heated to reflux for 16 hours. Upon completion, the reaction was cooled to room temperature, then the solvent was removed under vacuum, giving a crude oil. The product was precipitated by adding acetone in an ice bath, yielding a yellow solid. The crude product was purified by chromatographic column using ethyl acetate/petroleum ether in a 1:1 ratio as the mobile phase. The desired compound **21b** was obtained with an 81% yield.

^1^H NMR (400 MHz, DMSO-*d_6_*): *δ* 8.78 (d, *J* = 5.1 Hz, 1H), 8.47 (dd, *J* = 10.5, 2.7 Hz, 1H), 8.28 – 8.23 (m, 1H), 8.22 – 8.18 (m, 1H), 8.08 (d, *J* = 8.4 Hz, 1H), 7.89 – 7.84 (m, 1H), 7.72 – 7.64 (m, 2H), 6.93 (d, *J* = 5.0 Hz, 1H). ^19^F NMR (376 MHz, DMSO-*d_6_*): *δ* -126.59.

MS (ESI): m/z calcd for C_15_H_9_FN_2_O_3_ [M + H]^+^ 285.25; found 285.06.

*Synthesis of 3-fluoro-4-(quinolin-4-yloxy)aniline (****21c****).* To a solution of 4-(2-fluoro-4-nitrophenoxy)quinoline (**21b**, 9.05 mmol, 1 eq) in a mixture of ethyl acetate/acetic acid (15:1, 50 mL), palladium on activated charcoal (Pd 10%, 0.9 mmol, 0.1 eq) was added under a hydrogen atmosphere. The reaction was left under magnetic stirring for 48 hours at room temperature. The completion of the reaction was verified through ESI-MS and TLC (ethyl acetate/petroleum ether 4:1). The reaction mixture was treated by filtering the crude through celite and washing the residue with MeOH (3x15 mL). The crude was purified via column chromatography using ethyl acetate/petroleum ether in a 7:3 ratio as the mobile phase. A brownish solid was obtained (68% yield).

^1^H NMR (400 MHz, DMSO-*d_6_*): *δ* 8.66 (t, *J* = 4.7 Hz, 1H), 8.30 (dd, *J* = 8.4, 0.9 Hz, 1H), 7.99 (s, 1H), 7.82 – 7.77 (m, 1H), 7.65 – 7.60 (m, 1H), 7.08 (t, *J* = 9.0 Hz, 1H), 6.59 – 6.51 (m, 2H), 6.48 – 6.43 (m, 1H), 5.49 (s, 2H). ^19^F NMR (376 MHz, DMSO-*d_6_*): *δ* -130.75.

MS (ESI): m/z calcd for C_15_H_11_FN_2_O [M + H]^+^ 255.27; found 255.17.

*Synthesis of N-(3-fluoro-4-(quinolin-4-yloxy)phenyl)-3-((4-methylpyrimidin-2-yl)amino)benzamide (****21****).* To a solution of 3-((4-methylpyrimidin-2-yl)amino)benzoic acid **21a** (0.71 mmol, 1.1 eq) in DMF (5 mL), HATU (0.71 mmol, 1.1 eq), and DIPEA (0.71 mmol, 1.1 eq) were added at 0 °C. After 15 minutes, compound **21c** (0.65 mmol, 1 eq), previously dissolved in DMF, was added dropwise to the reaction mixture. The ice bath was removed, and the mixture was stirred at room temperature for 16 hours. Upon verifying the completion of the reaction through ESI-MS and TLC (ethyl acetate/petroleum ether 6:4), the solvent was removed under vacuum. The reaction mixture was extracted with ethyl acetate and washed with 10% citric acid, followed by 5% NaHCO_3_, and brine. The crude was purified via semi-preparative RP-HPLC using a gradient of acetonitrile/water (with 0.1% trifluoroacetic acid) at a flow rate of 20 mL/min. The desired product was obtained in a 21% yield.

^1^H NMR (400 MHz, DMSO-*d_6_*): *δ* 10.61 (s, 1H), 9.78 (s, 1H), 8.97 (d, *J* = 6.0 Hz, 1H), 8.53 (d, *J* = 8.4 Hz, 1H), 8.38 – 8.35 (m, 2H), 8.18 (d, *J* = 8.5 Hz, 1H), 8.15 – 8.04 (m, 2H), 8.00 (d, *J* = 8.0 Hz, 1H), 7.89 (t, *J* = 7.7 Hz, 1H), 7.75 (d, *J* = 8.9 Hz, 1H), 7.58 (t, *J* = 9.0 Hz, 1H), 7.53 – 7.39 (m, 2H), 7.00 (d, *J* = 5.9 Hz, 1H), 6.77 (d, *J* = 5.0 Hz, 1H), 2.37 (s, 3H). ^19^F NMR (376 MHz, DMSO-*d_6_*): *δ* -128.60, -128.62, -128.66. ^13^C NMR (DMSO-*d_6_*): *δ* 168.08, 166.91, 160.18, 157.99, 149.55, 141.50, 139.77, 135.60, 135.24, 133.90, 128.99, 124.41, 124.23, 122.77, 122.36, 120.51, 120.26, 118.54, 117.52, 109.44, 109.21, 104.28, 24.12.

HRMS (LC-MS Base Peak FullMS esi ^+^): m/z calcd for C_27_H_20_FN_5_O_2_ [M+H]^+^ 466.1674; found, 466.1655; error: -4.0328. T_R_ = 16.72 min (Fig. S2 and S3).

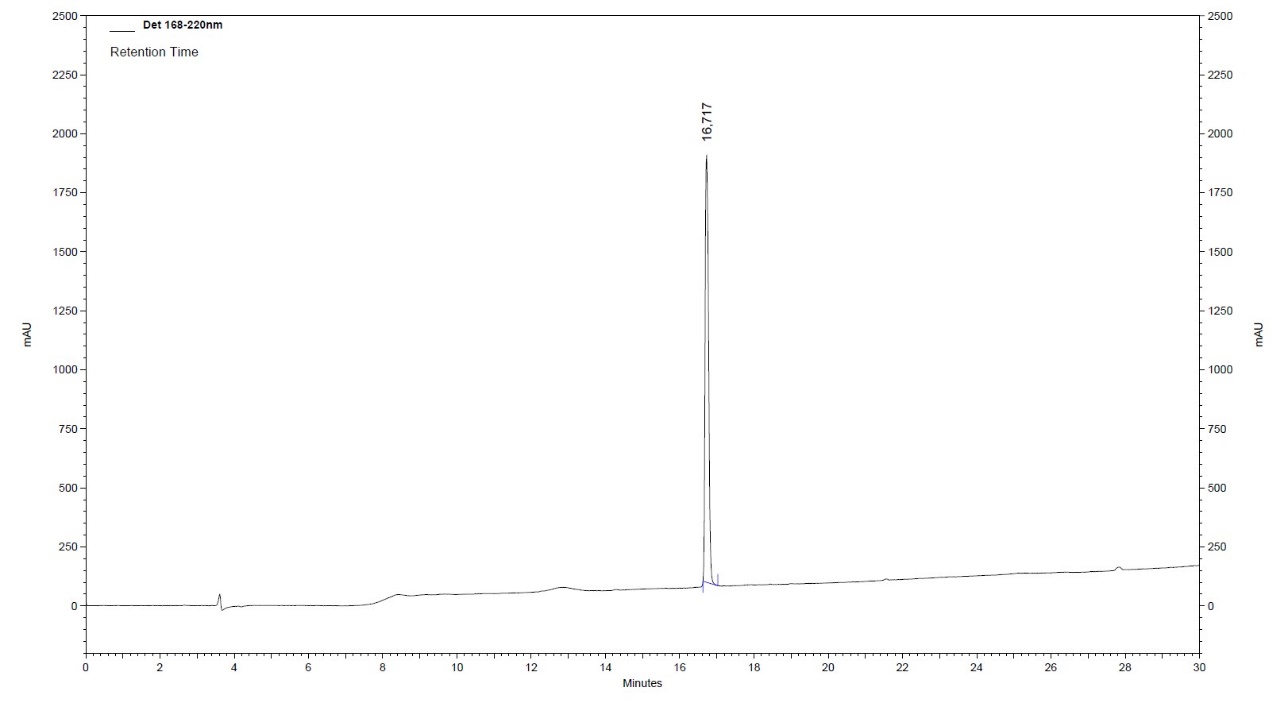

**Fig. S2.** Analytical HPLC profile of **21**.

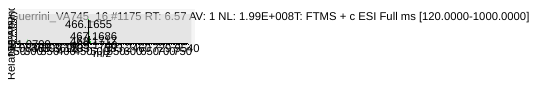

**Fig. S3**. High-resolution mass spectrum of final compound **21**.
